# Improving recombinant protein production using TISIGNER.com

**DOI:** 10.1101/2020.07.09.196469

**Authors:** Bikash K. Bhandari, Chun Shen Lim, Paul P. Gardner

## Abstract

Planning experiments using accurate prediction algorithms could mitigate failures in recombinant protein production. We have developed TISIGNER.com with the aim of addressing the technical challenges in recombinant protein production. We offer two web services, TIsigner (Translation Initiation coding region de<u>signer</u>) and SoDoPE (<u>So</u>luble <u>Do</u>main for <u>P</u>rotein <u>E</u>xpression), which are specialised in prediction/optimisation of recombinant protein expression and solubility, respectively. Importantly, TIsigner and SoDoPE are linked, which allows users to switch between the tools when optimising their genes of interest.

## INTRODUCTION

Recombinant protein production is the key process in supporting life science research and the development of biotherapeutics. However, low protein expression and solubility are the two major bottlenecks of recombinant protein production (1–5). Since mRNA abundance alone is insufficient to explain protein abundance at turnover (6–9), several features of mRNA sequence have been proposed to affect protein expression. These features are either related to codon usage, such as codon adaptation index and tRNA adaptation index (10–14), or mRNA secondary structure, such as G+C content, minimum free energy of RNA secondary structure, and mRNA:ncRNA interaction avoidance (15–20). Many of these features are not independent, making it hard to distinguish the impacts of individual features (21). This, in turn, hinders the development of accurate prediction/optimisation tools. However, recent systematic studies suggest mRNA secondary structure (minimum free energy) is the most important feature in protein expression (21, 22). We have recently shown that the mRNA accessibility of translation initiation sites outperforms mRNA secondary structure (minimum free energy) in predicting the expression of 11,430 recombinant proteins (23).

In addition to high expression level, high solubility is preferable for the purification and long-term storage of recombinant proteins. However, almost half of the successfully expressed proteins are insoluble (http://targetdb.rcsb.org/metrics/), which makes the recombinant protein production process more challenging. A number of methods have been suggested to improve protein solubility, for example, truncation, mutagenesis, and the use of solubility-enhancing tags (1, 24–26). Nevertheless, accurate solubility prediction could save resources and aid in designing soluble proteins before the experiment. With these in mind, we have recently formulated the Solubility-Weighted Index (SWI), which outperforms recent solubility prediction tools based on machine-learning algorithms (27).

Many existing tools predict or optimise either protein expression or solubility alone. We reasoned these functionalities should be integrated as high levels of recombinant protein expression and solubility are both desirable in most downstream applications. Here we present TISIGNER.com that integrates the optimisation tools TIsigner (<u>T</u>ranslation <u>I</u>nitiation coding region de<u>signer</u>) and SoDoPE (<u>So</u>luble <u>Do</u>main for <u>P</u>rotein <u>E</u>xpression) for protein expression and solubility, respectively. Our web application provides easy, fast and interactive ways to assist users in planning and designing their experiments.

## WEB SERVICES

### TIsigner

TIsigner offers tunable protein expression by optimising the mRNA accessibility of translation initiation sites (23). The regions used to calculate accessibility (opening energy) are specific to the expression hosts, which is calculated using RNAplfold (28–30). For *Escherichia coli*, *Saccharomyces cerevisiae*, and *Mus musculus* expression hosts, the regions for optimisation are −24:24, −7:89, −8:11 respectively. For other expression hosts, we provide an option ‘Other’, which optimises the accessibility of the region −24:89. Since *E. coli* is the most popular expression host, the default settings aim to optimise protein expression in the *E. coli* T7 lac promoter system (see below). In this case, only the protein coding sequence is needed as input (Fig 1). Otherwise, the 5′ UTR (5′ untranslated region) sequence is also required as input.

**Fig 1.**
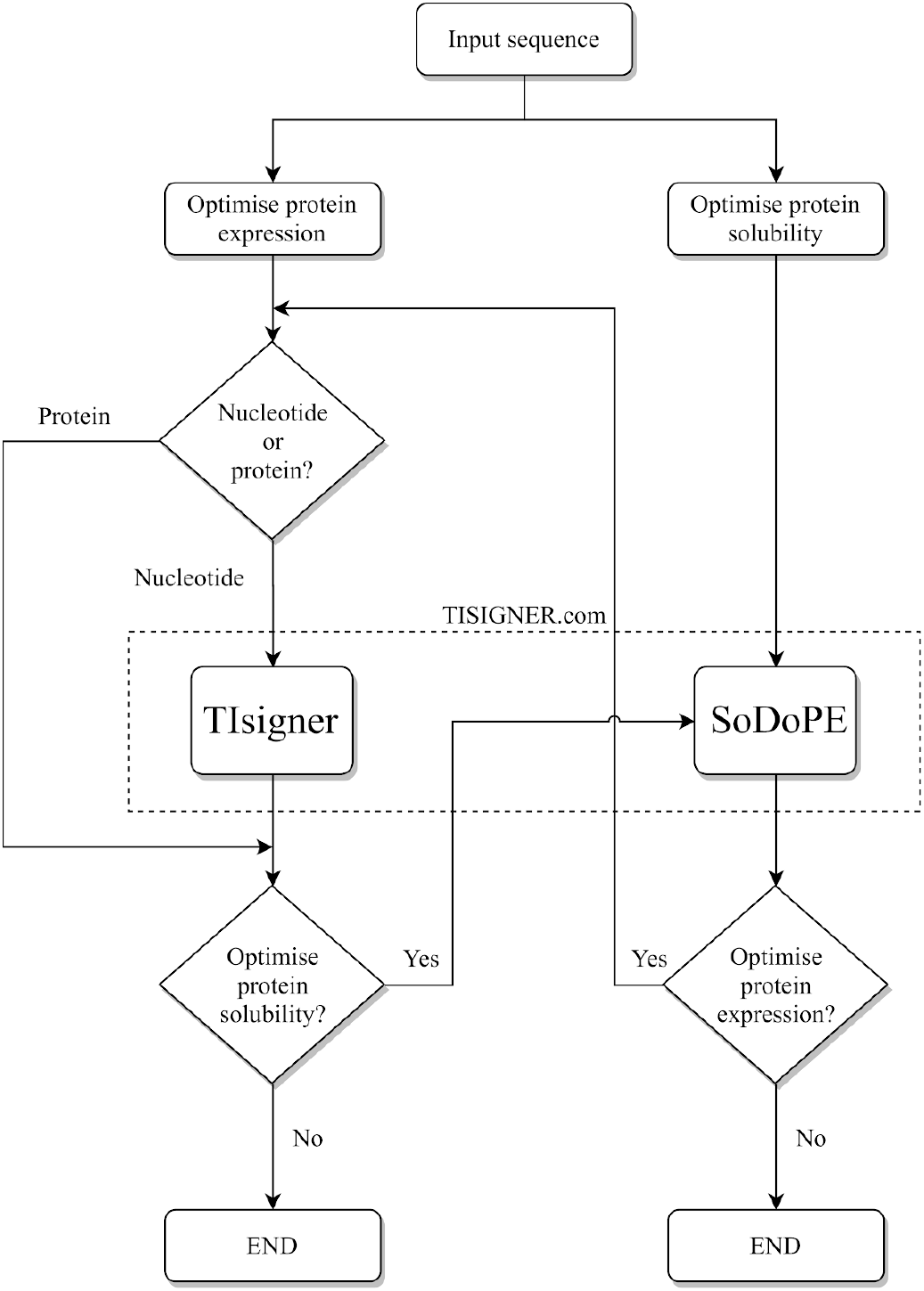
Flow chart for optimising recombinant protein production using the TISIGNER web application. TIsigner and SoDoPE are linked so that protein expression and solubility can be seamlessly optimised. TIsigner accepts a nucleotide sequence as input, whereas SoDoPE accepts either a nucleotide or protein sequence. SoDoPE, <u>So</u>luble <u>Do</u>main for <u>P</u>rotein <u>E</u>xpression; TIsigner, <u>T</u>ranslation <u>I</u>nitiation coding region de<u>signer</u>.

The settings for TIsigner are grouped by complexity (i.e., general, extra and advanced). The general settings include the options to modify the expression host, promoter and target expression score. The target expression score is only applicable to the *E. coli* T7 lac promoter system as the score is calculated based on the opening energy distribution of 11,430 expression experiments in *E. coli* from the ‘Protein Structure Initiative:Biology’ (PSI:Biology) (31, 32). For other expression hosts and promoters, the target expression level can be either maximised or minimised (i.e., binary). The extra settings have the options to optimise sequence within the translation initiation region or full-length sequence. The AarI, BsaI, BsmBI restriction modification sites are filtered by default, whereas other sites can be manually supplied. The advanced settings allows users to tweak the random seed and sampling options (i.e., quick or deep, which uses different numbers of iterations and parallel processes). Here users can also customise the region for optimisation.

Once the input sequence passes through a sanity check, the optimisation task takes only O(1) time using RNAplfold (version 2.4.11; using parameters -W 210 -u 210) with a simulated annealing algorithm. A list of optimised sequences will be returned after checking for terminators using cmsearch (INFERNAL version 1.1.2) (33) and RMfam (34, 35). If terminators are found, an option to use the full-length sequence for optimisation will be prompted to users. In a default case (*E. coli* T7 lac promoter system), the optimised sequence closest to the chosen expression level will be shown as the first solution. For other expression hosts and/or promoters, the optimised sequence with the minimum number of mismatches will be shown as the first solution. These mismatched nucleotides will be highlighted. The accessibility of translation initiation sites for both the input and optimised sequences will be shown as opening energy (kcal/mol). These results can be exported as a PDF or CSV file. When the default settings are used, the opening energy for each sequence will be indicated on the distributions of the opening energy of 8,780 ‘success’ and 2,650 ‘failure’ groups of the PSI:Biology target genes. Furthermore, an option for solubility analysis using SoDoPE (see below) will be available for each sequence on the same results page.

### SoDoPE

SoDoPE is our interactive solubility analysis and optimisation tool based on SWI (27). SoDoPE accepts either a nucleotide or protein sequence (Fig 1). Upon sequence submission, a query will be sent to the HMMER web service for protein domain annotation (36). Successful annotation will be displayed as interactive graphics, in which the annotated domains will be represented as stadiums that overlay a grey band representing the input protein sequence. Information about a protein domain will be shown upon a mouse hover (Fig 2). These domains can be selected for solubility analysis. For a complete domain annotation report, a link to the HMMER results page will be provided.

**Fig 2.**
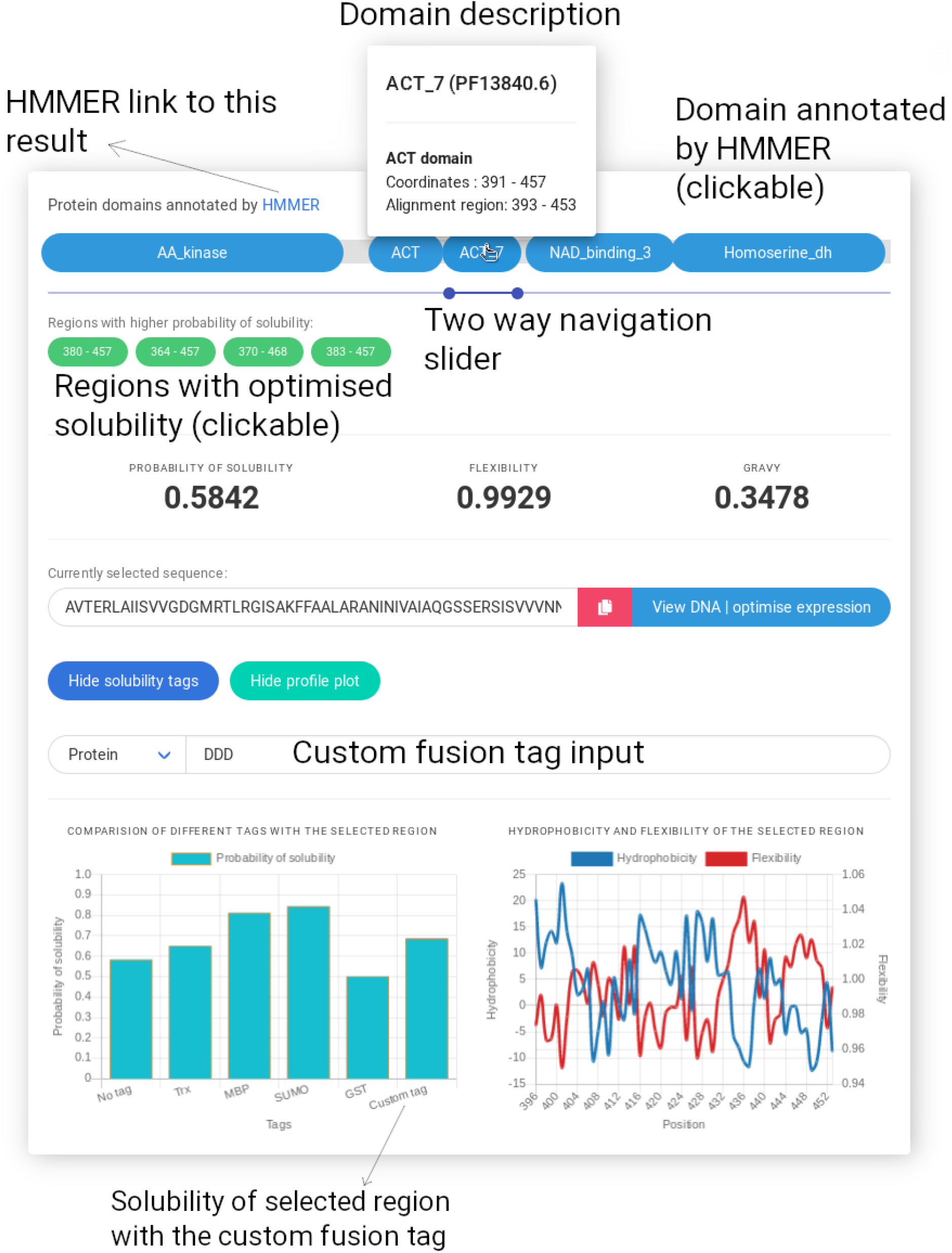
Exploring and optimising protein solubility using SoDoPE interactive graphics. Upon clicking a protein domain or selecting a region of interest, its solubility will be optimised in real-time, and a list of regions with extended boundaries and higher probabilities of solubility will be returned as green buttons (clickable). The probabilities of solubility of the selected region with and without fusion tags can be visualised in a barplot. The flexibility and hydrophobicity profile plots for the selected region can also be selectively viewed.

In addition, a two-way slider will be available for navigation through any region of interest. The probability of solubility, flexibility and GRAVY (grand average of hydropathy) will be shown in real-time according to the user-selected region. The selected region will be optimised for higher solubility using simulated annealing. Only the regions with extended boundaries and also higher probability of solubility will be returned.

A profile plot of flexibility and/or hydrophilicity corresponding to the user selected region can be generated. This allows an estimation of rigid/flexible regions and possible helices, which might be helpful for mutagenesis experiments. The sequence of the selected region will be shown, with the option of sequence conversion between nucleotide and amino acid sequence format. In particular, the nucleotide sequence can be redirected to TIsigner for optimisation of protein expression.

The contributions of several solubility-enhancing tags to the user selected region can be compared and shown in a bar plot, including thioredoxin (TRX), maltose binding protein (MBP), small ubiquitin-related modifier (SUMO) and glutathione S-transferase (GST) tags. Users can also input a fusion sequence of interest either in a nucleotide or protein sequence format.

## GENERAL INFORMATION

Demo input and results are available for new users to get started. A list of frequently asked questions is also available for each tool. The frontend is written in React and uses responsive web design principles. The backend is written in Flask and Python 3.6. The website is hosted on a virtual machine (Red Hat Enterprise Linux 8) running on Intel Xeon (8 × 2.60 GHz) with 4GiB RAM, by the Information Technology Services at the University of Otago.

## ACKNOWLEDGEMENTS

This work was supported by the Ministry of Business, Innovation and Employment, New Zealand (MBIE grant: UOOX1709).

